# Reconstruction of Gene Regulatory Networks using sparse graph recovery models

**DOI:** 10.1101/2023.04.02.535294

**Authors:** Harsh Shrivastava

## Abstract

There is a considerable body of work in the field of computer science on the topic of sparse graph recovery, particularly with regards to the innovative deep learning approaches that have been recently introduced. Despite this abundance of research, however, these methods are often not applied to the recovery of Gene Regulatory Networks (GRNs). This work aims to initiate this trend by highlighting the potential benefits of using these computational techniques in the recovery of GRNs from single cell RNA sequencing or bulk sequencing based gene expression data. GRNs are directed graphs that capture the direct dependence between transcription factors (TFs) and their target genes. Understanding these interactions is vital for studying the mechanisms in cell differentiation, growth and development. We categorize graph recovery methods into four main types based on the underlying formulations: Regression-based, Graphical Lasso, Markov Networks and Directed Acyclic Graphs. We selected representative methods from each category and made modifications to incorporate transcription factor information as a prior to ensure successful reconstruction of GRNs.

## 1 Introduction

In molecular biology, it is well understood that the expression levels of genes are controlled by transcription factors (TFs). These TFs are proteins that regulate the expression levels of their target genes in a specific cell at a given time. These regulatory relationships can be represented by a graph, called a gene regulatory network (GRN), where nodes represent genes, and an edge from gene A to gene B indicates that the protein product of gene A is a TF that regulates gene B. This network governs transcription and ultimately determines how cells behave, making it a topic of great interest to scientists who aim to decipher the interactions within this network. It has been a long-standing challenge to reconstruct these networks computationally from gene-expression data [6, 19]. Recently, developed single cell RNA-Sequencing (scRNA-Seq) technologies provide unprecedented scale of genome-wide gene-expression data from thousands of single cells. This access to large amounts of higher quality expression data should be leveraged to the inference of more reliable and detailed regulatory networks. Some works [4, 25] have already bench-marked existing GRN recovery methods.

In this work, we will start by briefly describing the popular state-of-the-art techniques for reconstructing Gene Regulatory Networks. We will then delve into the broader category of graph recovery methods, many of which have been developed for purposes other than computational biology. Additionally, we will highlight some of the most recent and effective deep learning based approaches. We provide essential modifications for some of the selective approaches to make them suitable for GRN reconstruction. We will provide links to various implementations and offer guidance on the general approach and considerations to keep in mind when adapting a graph recovery method for GRN reconstruction. It is our hope that this overview will give the reader a comprehensive understanding of the various graph recovery methods and empower them to apply these techniques to the task of GRN reconstruction. We believe that this information will be valuable for researchers and practitioners alike. So, the reader will be able to make informed decisions when selecting the appropriate method for their specific needs.

## 2 Overview of Graph Recovery Approaches

### 2.1 Problem setting

We consider gene expression data as input *X* ∈ *R*^*M*×*D*^, which consists of *D* genes and *M* samples. Let 𝒢= [1, ⋯, *D*] be the set of genes and 𝒯 ⊂ 𝒢denote the subset of genes that are Transcription Factors. We assume that we already know the list of TFs for a given input. Our aim is to identify the directed interactions of the form (*t, o*), where *t* ∈ 𝒯 and *o* ∈ 𝒢. We acknowledge that there may be interactions between the TFs. However, for the purposes of the methods discussed in this work, we choose to disregard these interactions. It is possible to include these interactions by assigning undirected edges between the connections found among the TFs. In these situations, the outcome will be Completed Partially Directed Acyclic Graphs (CPDAGs), where PDAGs represent equivalence classes of Directed Acyclic Graphs and stands for ‘partially directed acyclic graphs’ [8].

Fig. 1 attempts to list popular formulations of graph representations and the corresponding algorithms to recover them. Modifications to include TF information for GRN reconstruction for the methods highlighted in blue will be discussed in detail in this work. In order to enhance the reader’s understanding of this field, we will briefly describe approaches in each of the categories outlined.

**Figure 1:**
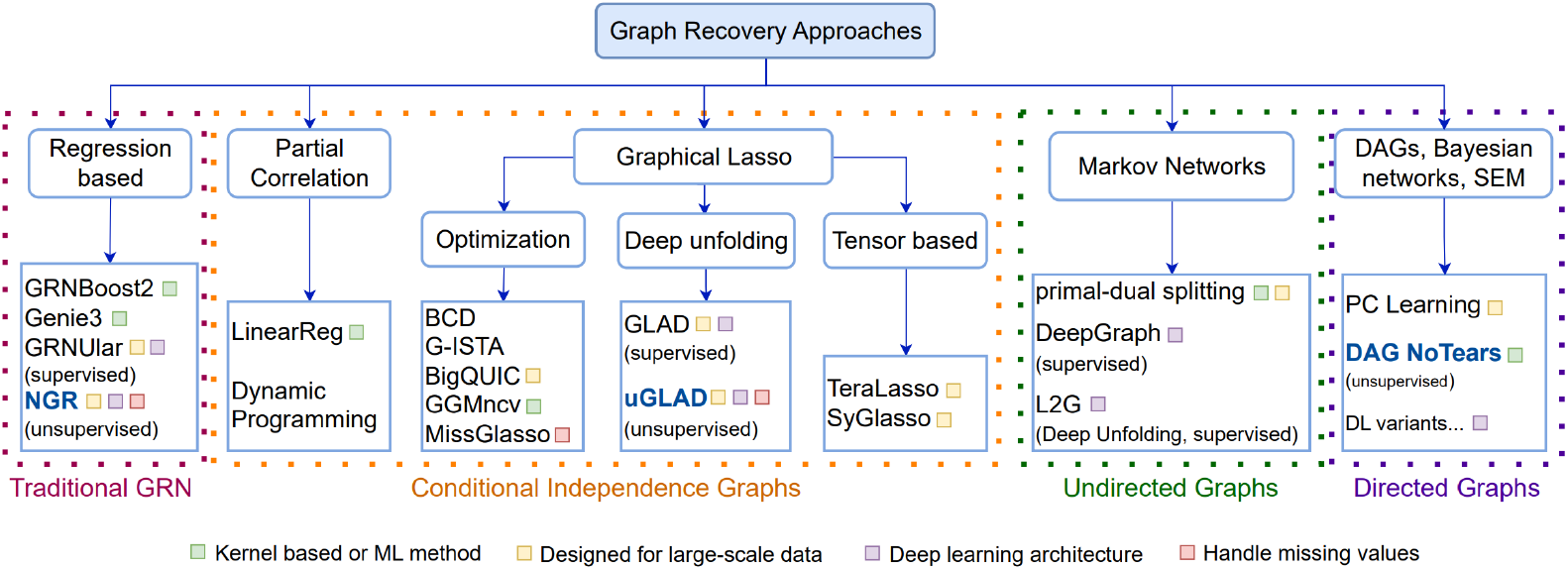
Graph Recovery approaches. Methods used to recover sparse graphs are categorized into four types based on the underlying formulation. The existing GRN recovery approaches mostly follow the regression based formulation (leftmost branch). The algorithms (leaf nodes) listed here are representative of the sub-category and the list is not exhaustive. Modifications for the highlighted methods are worked out in this paper to include the TF information for the GRN recovery task.

### 2.2 Regression based approaches for GRN Reconstruction

Many of the state-of-the-art methods that are traditionally used for inferring gene regulatory networks involve fitting regression functions between the expression values of transcription factors and other genes that may be potential targets. A sparsity constraint is often applied to the regression function in order to identify the most influential TFs for each gene. The objective function used for GRN recovery in various methods is typically a variant of the equation provided below

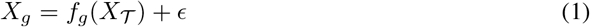

We can interpret Eq. 1 as fitting a regression between each gene’s expression value and the expression values of TFs, with some added random noise. A simple model for this would be to assume that the function fg is linear. One method, TIGRESS [14], uses a linear function of the following form for each gene, *f*_*g*_ (*X* _𝒯_) = Σ_*t* ∈ 𝒯_ *β*_*t,g*_ *X*_*t*_. Another top performing method GENIE3 [39], assumes each *f*_*g*_ to be a random forest. GRNBoost2 [23] further uses gradient boosting techniques over the GENIE3 architecture to do efficient GRN reconstruction. Recently developed deep unfolding based deep learning methods like GRNUlar [32, 35] became the new state-of-the-art models. Ensemble methods like ‘EnGRaiN’ [1] also enhances the performance of the traditional approaches. In this paper, we additionally explore a new type of probabilistic models known as Neural Graphical Models (NGMs) [29] used neural network as a multitask learning to fit regressions and recover graph for generic input datatypes. We specifically focus on ‘Neural Graph Revealers’ (NGRs) [30] architecture and re-purpose it for the GRN reconstruction task.

### 2.3 Conditional independence graphs

conditional independence (CI) graphs as undirected probabilistic graphical models that show partial correlations between features. CI graphs are primarily used to gain insights about feature relationships within a system to help with decision making ans system designing. In some cases, they are also used to study the evolving feature relationships with time [18]. Given *m* observations of a *d*-dimensional multivariate Gaussian random variable *X* = [*X*_1_, ⋯, *X*_*d*_]^⊤^, the sparse graph recovery problem aims to estimate its covariance matrix Σ^*^ and precision matrix Θ^*^ = (Σ^*^)^−1^. The *ij*-th component of Θ^*^ is zero if and only if *X*_*i*_ and *X*_*j*_ are conditionally independent given the other variables {*X*_*k*_} _*k* ≠*i,j*_. The general form of the graphical lasso optimization to estimate Θ^*^ is the log-likelihood of a multivariate Gaussian with regularization as

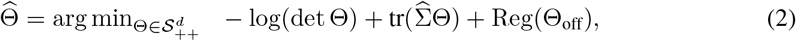

where 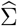 is the empirical covariance matrix based on *m* samples, 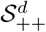 is the space of *d* × *d* symmetric positive definite matrices and Reg(Θ_off_) is the regularization term for the off-diagonal elements. Once the precision matrix is obtained, the corresponding partial correlation matrix entries can be calculated as 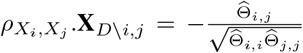 (definition taken from [28]). Several algorithms have been developed to optimize the sparse precision matrix estimation problem in Eq. 2 which primarily differ in the choice of regularization and optimization procedure. A survey on ‘Methods for recovering Conditional Independence graphs’ [28] and other works like [11, 17, 9, 41, 3, 12, 42] extensively discuss the different formulations and applications for methods in this category. We choose the deep unfolding (or unrolled algorithm) based model `GLAD` [31] and its unsupervised version `uGLAD` [33, 34] to extend for GRN reconstruction.

### 2.4 Markov Networks

Markov networks are probabilistic graphical models defined on undirected graphs that follow the Markov properties [20]. While Markov networks follow the conditional independence properties which can be pairwise, local and global, we made the distinction from the Conditional Independence graphs based on the interpretation of the edges. They can be learned using traditional constraint-based and score-based structure learning methods, but these methods can suffer from combinatorial explosion of computation requirements and often require simplifying approximations. Recently, researchers have developed supervised deep learning architectures, such as ‘DeepGraph’ [2] to learn mapping from samples to a graph. However, these methods have a large number of learning parameters and have shown limited success. An alternative approach is the “deep unfolding” or “unrolled algorithm” methodology, which were primarily developed for compressed sensing have now been used to recover Markov networks such as Iterative Shrikage Thresholding Algorithm (ISTA) [13], ALISTA [22] and others [37, 7, 5]. One example of this approach to learn graph topology, is the L2G framework, which unrolls a primal-dual splitting algorithm into a neural network for their model [26].

### 2.5 Directed Graphs

This is an active area of research with many new methods being developed quickly. There are several types of directed graphs that are commonly studied, such as Directed Acyclic Graphs, Bayesian Networks, and Structural Equation models. One of the first techniques developed to learn Bayesian Networks was the PC-learning algorithm [36], followed by a suite of score-based and constraint-based learning algorithms and their parallel variants [15, 20]. Recently, a new method called ‘DAG with NOTEARS’ [45] was introduced which converts the combinatorial optimization problem of DAG learning into a continuous one. This has led to the development of many follow up works, including some deep learning methods [43, 44, 46, 24]. There are also many reviews of causal structure learning methods and structural equation models available, for instance the work in [16]. In this work, we propose modifications to the “DAG with NOTEARS” formulation to incorporate the TF prior for GRN reconstruction.

## 3 Newer methods for GRN Reconstruction

We kindly assume that our readers are already familiar with the original formulations of the methods discussed below, refer literature provided in Sec. 2. In order to set the stage for introducing the modifications, we will only include the basic notation and definitions that are necessary. Table 1 provides the links to the implementations of the discussed methods.

**Table 1:** sparse graph recovery methods with their implementation links. Additional information about their ability to recover multiple graphs (MG), handle large scale data (LS) and handle missing values in data (MI) are mentioned alongside.

| Methods | Implementation | MG | LS | MI | Paper |
| --- | --- | --- | --- | --- | --- |
| DAG NoTears | https://github.com/xunzheng/notears |  | ✓ |  | [45] |
| uGLAD | https://github.com/Harshs27/uGLAD | ✓ | ✓ | ✓ | [34] |
| NGR | https://github.com/harshs27/neural-graph-revealers |  | ✓ |  | [29] |

### 3.1 Neural graph revealers for GRNS

#### Representation

We consider the input expression data *X* ∈ R^*M* ×*D*^, with *D* = {*T* ∪ *G*} genes and *M* samples. The task is to recover a sparse graph represented by a dependency matrix of the form *S*_*G*_ ∈ R^*T* ×*G*^. The architecture of NGR is a multilayer perceptron (MLP) that takes in input TFs and fits a regression to get the target genes as the output, shown in Fig. 2(left). We view this neural network as a glass-box where a path to an output unit (or neuron) from a set of input units means that the output unit is a function of those input units. In order to obtain a GRN, we want to find the most relevant TFs that have a direct functional influence for every target gene *g*_*d*_. This task becomes increasingly complex as we need to evaluate all possible combinations which can be computationally tedious. Fitting of regression of NGRs, refer Fig. 2(middle), can be seen as doing *multitask learning* that simultaneously optimizes for the functional dependencies of all the TF and target genes. We now want to efficiently induce sparsity among the paths defined by the MLP.

**Figure 2:**
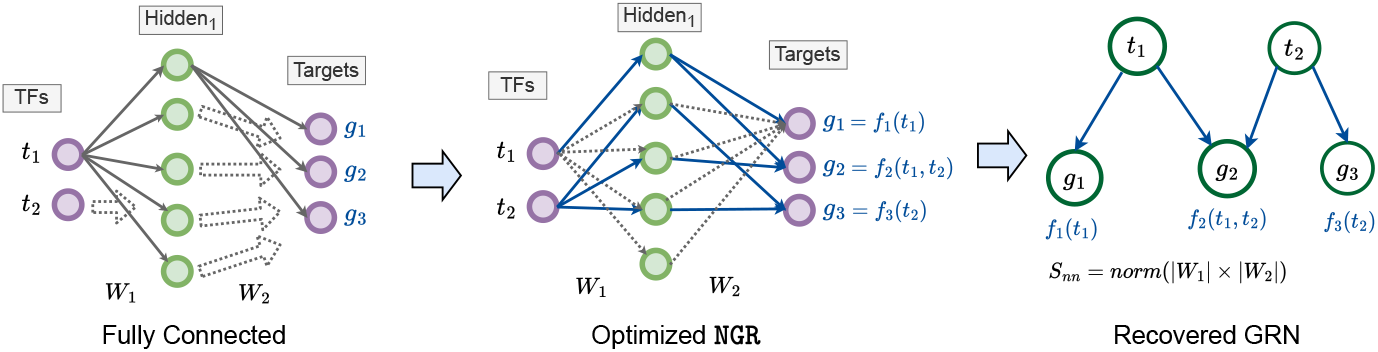
Modified workflow of NGRs for GRN recovery. (left) We start with a fully connected Neural Network (MLP here) where both the input are the TFs and the output are the target genes. Viewing NN as a multitask learning framework indicates that the output genes are dependent on all the input TFs in the initial fully connected setting. (middle) The learned NGR optimizes the network connections to fit the regression as well as satisfy the sparsity constraints, refer Eq. 3. If there is a path from the input TF to an output gene, that indicates a potential non-linear dependency between them. By varying the size of NN (number of layers, hidden unit dimensions), the complexity of the functional representation can be controlled. Note that not all the weights of the MLP (those dropped during training in grey-dashed lines) are shown for the sake of clarity. (right) The sparse dependency graph (bipartite) between the input and output of the MLP reduces to its normalized weight matrix product *S*_*nn*_ = norm (|*W*_1_| *×* |*W*_2_|).

##### Algorithm 1: Recovering GRNs using NGRs

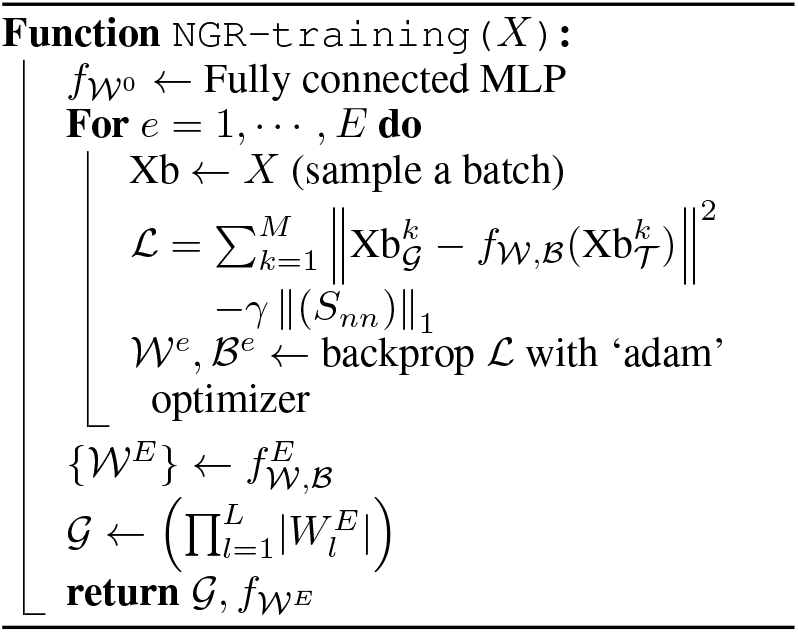

#### Optimization

(notations adapted from the NGR paper [29]). We denote a NN with *L* number of layers with the weights 𝒲 = {*W*_1_, *W*_2_, ⋯, *W*_*L*_} and biases ℬ = {*b*_1_, *b*_2_, ⋯, *b*_*L*_} as *f* _𝒲,ℬ_ (·) with non-linearity as ReLU. Applying the NN to the input *X* evaluates the following mathematical expression, *f* _𝒲, ℬ_ (*X*) = ReLU(*W*_*L*_ · (⋯ (*W*_2_·ReLU(*W*_1_ · *X* + *b*_1_) + *b*_2_) ⋯) + *b*_*L*_). The dimensions of the weights and biases are chosen such that the neural network input units are equal to 𝒯 and output units are equal to 𝒢, while the hidden layers dimension *H* remains a design choice that is adjusted based on the regression results on the validation data. We note that if *S*_*nn*_[*t*_*i*_, *g*_*o*_] = 0 then the target gene *g*_*o*_ is not influenced by the input TF *t*_*i*_. We use these formulations of MLPs to model the constraints along with finding the set of parameters { 𝒲 ℬ }, that minimize the regression loss expressed as the Euclidean distance between *X*_𝒢_ to *f* _𝒲,ℬ_, (*X* _𝒯_). We induce sparsity using the *ℓ*_1_ norm as a Lagrangian term, thus the optimization becomes

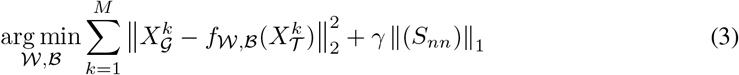

where the product of the weights of the neural networks 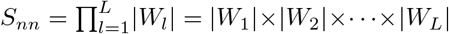 gives us path dependencies between the input TFs and the output target genes. We can optionally add log scaling to the sparsity constraint term for more stable convergence. Essentially, we start with a fully connected graph and then the Lagrangian term induces sparsity in the GRN. Alg. 1 describes the procedure to optimize the modified NGR architecture based on Eq. 3. The optimization and the graph recovered depend on the choice of the penalty constant gamma. To get a good initial value of the constants, the loss-balancing technique introduced in [27] can be utilized.

Interestingly, this modification of NGR can also be seen as extending the `GRNUlar` [35] framework into an unsupervised GRN reconstruction model.

### 3.2 UGLAD FOR GRNS

We modified the `uGLAD` algorithm [33], refer Alg. 2, to take into account TF information, called `uGLAD`GRN, by using a post-hoc masking operation that only retains the edges having at least one node as a transcription factor. The `GLAD` cell utilized as a subroutine of the algorithm is available in Alg.1 of their paper. This post-hoc operation is very generic and can be applied to most of the algorithms that recover Conditional Independence graphs.

#### Algorithm 2: uGLAD-GRN

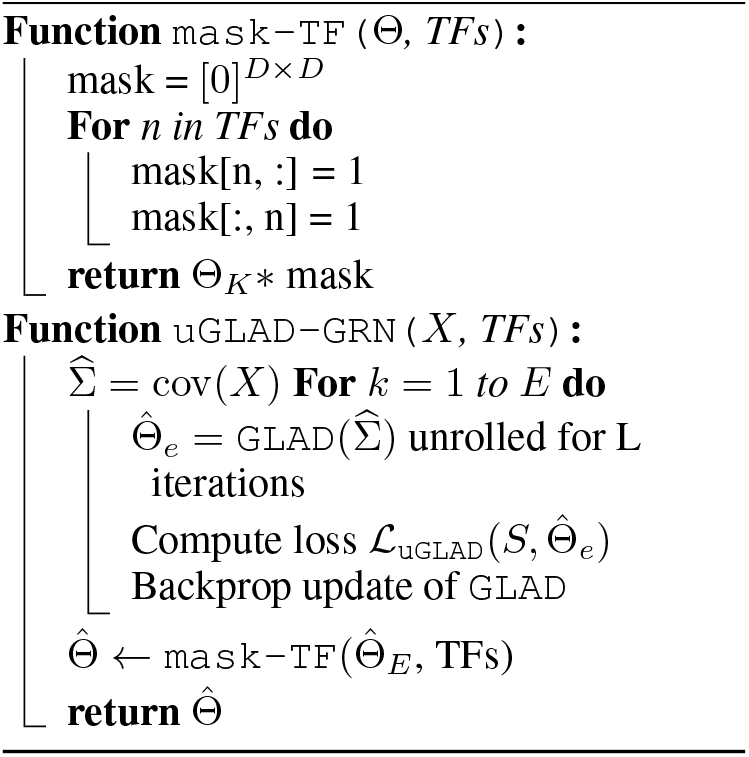

### 3.3 DAG With NOTEARS For GRNS

We follow the notations as defined in their paper [45], for the ease of the readers. We refer to Eq.10 of their paper and directly add the changes needed in the optimization to solve the equality-constrained program for linear Structural Equation Model (SEM) below

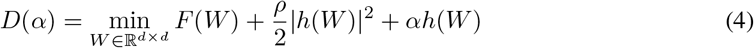

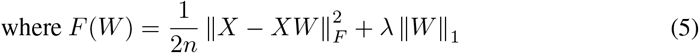

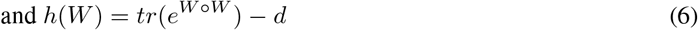

We will make modifications in the *F* (*W*) term to account for the TFs. Specifically, our changes ensure that the recovered GRN will have no incoming edges to the TFs.

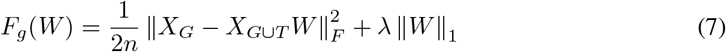

where, *X*_*G*_ contains data only for the target genes and total genes *D* = { *G* ∪ *T* }. The rest of the details about optimization are the same as in the Eq.11 and Eq.12 of the paper which utilizes the augmented Lagrangian method. We just replace the *F* (*W*) with the *F*_*g*_(*W*). Note that we do not obtain a complete bipartite graph in this case as there can be directed edges between the genes.

### 3.4 Using Biological data SIMULATORS

While the techniques for GRN reconstruction from expression data are improving, simulators which generate synthetic expression data guided by GRNs are also advancing [40, 25, 21]. These biological gene expression simulators can generate realistic data, and have modeled sources of variation such as noise intrinsic to transcription, extrinsic variation indicative of different cell states, technical variation, and measurement noise and bias. These realistic simulators are primarily used to benchmark the performance of GRN inference methods [10, 4, 25]. Evaluations of current methods for GRN inference show that their performance is not satisfactory, even with synthetic data. GRN inference is hindered by multiple factors, such as the potentially nonlinear relationships between transcription factors (TFs) and their target genes, and the intrinsic and technical noise present in expression data including the single-cell RNA sequencing data [38]. SERGIO [10] is a realistic GRN-guided simulator for scRNA-Seq data, which has been used to train and evaluate GRN recovery models. It generates realistic gene expression data by incorporating known principles of TF-gene regulatory interactions that underlie expression dynamics, and models the stochastic nature of transcription as well as simulating the non-linear influences of multiple TFs.

The works [31, 32] were one of the first models that utilized the data from such biological simulators, SynTReN [40] & SERGIO [10] respectively, for training and evaluation. They posit that the models can learn to capture the underlying distribution from which the GRNs and their corresponding expression data are sampled by utilizing various simulator settings. They demonstrate with varied level of success that their models were able to transfer their learning to recover GRNs on real experimental data. We plan to release extensive evaluations of these approaches on the simulated as well as on the real expression data soon.

## 4 Conclusions

Reconstruction of Gene Regulatory Networks is an important problem from biological insights perspective and also pose interesting computational challenges with newer sequencing technologies that are being developed at an unprecedented pace. Our research project aims to bridge the gap between computer science and biology by exploring the potential of sparse graph recovery algorithms for reconstructing Gene Regulatory Networks from gene expression data. To do this, we have classified the sparse graph recovery approaches and identified recent and representative unsupervised deep learning models from these categories. We have also worked out the modifications needed to make these approaches suitable for the GRN reconstruction task and provided links to their implementations.

